# Trimodal brain-wide ultrasound imaging of brain-tumor interaction

**DOI:** 10.1101/2025.10.29.685462

**Authors:** Claire Rabut, Shirin Shivaei, Baptiste Heiles, Mikhail G. Shapiro

## Abstract

Patients with brain tumors often suffer debilitating neurological dysfunction as their tumors disrupt brain tissue, affecting both local and global neural activity and blood flow. However, studying tumor-brain interactions in animal models is challenging due to a lack of methods that simultaneously capture dynamics of tumor growth, neural activity and vascular alterations over time. Here, we overcome this limitation using a multimodal ultrasound imaging platform, an imaging technique that offers brain-wide coverage in living animals at 10-100 µm resolution. To monitor the co-evolution of tumor growth, neural activity and vascular remodeling, we integrated (1) nonlinear imaging of acoustic reporter gene expressing-tumors, (2) hemodynamic functional imaging of brain activity, and (3) super-resolution microscopy of the vasculature. Integrating these modalities for the first time and applying them to a common model of glioblastoma, we followed tumor-brain interactions in individual animals over their disease lifetimes. Our approach allowed us to precisely map the spatial displacement of functional brain regions, the local and global disruption of functional connectivity, and the remodeling of the blood supply to support tumor growth. This integrated method bridges a critical gap in brain cancer research and therapy development by providing a unified dynamic view of what happens to the brain as tumors grow within it.

## INTRODUCTION

The interaction between brain tumors, such as gliomas, and native cells in the brain can reshape whole-brain activity, vasculature and metabolism. Previous studies have shown significant changes in neural activity patterns in both near and distant sites from brain tumors^1^. Brain tumors also remodel the brain vasculature by inducing the growth of new leaky and dysfunctional blood vessels^2^, compromising the blood-brain barrier, causing edema and increasing intracranial pressure^3^. Furthermore, the competition between tumor cells and neurons for metabolic resources can exacerbate neuronal dysfunction and degeneration^4^. Together, these mechanisms of disruption can cause profound effects on brain health, function and patient well-being^5^. Several imaging modalities have been applied to brain–tumor studies, but each faces critical limitations. MRI provides whole-brain coverage and is widely used to track tumor growth and vasculature^6^, yet it doesn’t allow awake behaving studies. Two-photon and wide-field optical methods reveal cellular or vascular dynamics with high resolution, but are limited to superficial cortex^7^. PET sensitively reports on metabolism, but with low spatiotemporal resolution and radiation constraints^8^. No existing method can non-invasively capture tumor growth, vascular remodeling, and functional neural activity together with high resolution and depth, motivating our development of a trimodal ultrasound platform.

Here, we tackle this challenge by developing a trimodal ultrasound imaging platform capable of simultaneously imaging tumors, vessels, and neural activity in behaving animals (**Fig. 1**). Ultrasound is uniquely suited to this task because it is portable, non-invasive, penetrates opaque tissue at centimeter depths, and provides high spatial resolution (∼100 µm). Recent advances have further expanded its capabilities into complementary domains relevant to tumor biology. Acoustic reporter genes (ARGs) based on gas vesicles (GVs), air-filled protein nanostructures derived from bacteria^9^, enable direct ultrasound visualization of genetically defined tumor cells, allowing longitudinal tracking of tumor growth with molecular specificity^10–13^. Functional ultrasound imaging (fUSI) leverages ultrafast Doppler sequences to map whole-brain hemodynamic activity in awake, behaving animals with high spatiotemporal resolution^14–16^. Finally, ultrasound localization microscopy (ULM) reconstructs microvascular networks by tracking circulating microbubbles at ∼10 µm resolution, breaking the diffraction limit by an order of magnitude^17^ and revealing tumor-driven angiogenesis and vascular remodeling^18–20^. Together, these complementary approaches position ultrasound as a uniquely versatile modality for simultaneous molecular, vascular, and functional imaging of brain–tumor interactions in individual animals. In this study, we combine ARGs, fUSI, and ULM to form a synergistic imaging platform to study the molecular, functional, and structural dynamics of brain–tumor interactions.

**Figure 1:**
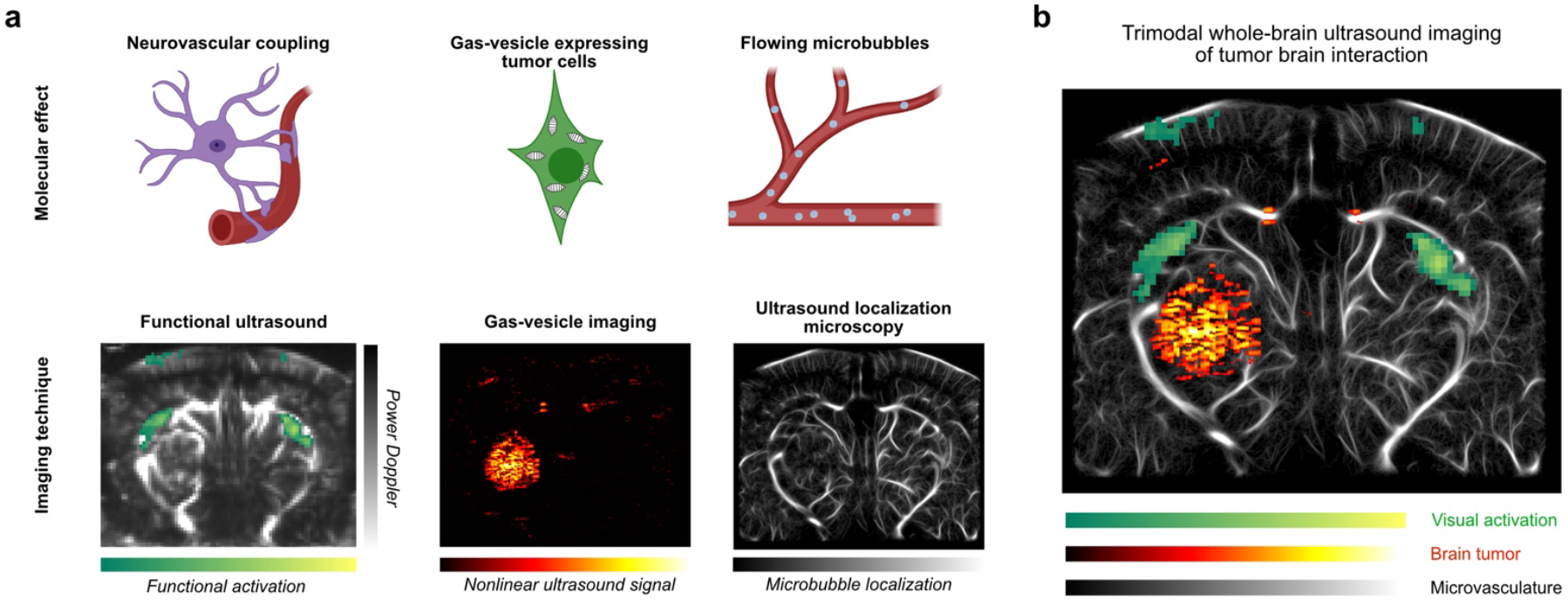
Trimodal ultrasound imaging platform for studying tumor–brain interactions. **a**. Schematic overview of the molecular effectors and corresponding imaging modalities used to capture tumor–brain interactions. Fron left to right: fUSI detects changes in cerebral blood flow linked to neurovascular coupling. GV imaging enables specific visualization of genetically encoded tumor cells via nonlinear contrast. Circulating microbubbles allow for high-resolution mapping of the microvascular network using ULM. **b**. Example of whole-brain trimodal imaging in a tumor-bearing mouse. The image shows spatially aligned maps of visual activation (green), GV-expressing tumor signal (hot colors), and microvascular structure (grey), illustrating the ability of the platform to simultaneously track neural activity, tumor progression, and vascular remodeling in vivo.

We demonstrate the capabilities of this platform by creating a stable GV-expressing glioblastoma cell line, implanting it in immunodeficient mice, and precisely mapping tumor growth and its consequences for sensory-elicited neural activity, functional connectivity and vascular remodeling in individual behaving animals. All three types of data are acquired during imaging sessions lasting less than one hour and repeated throughout the tumor lifetime of the animal. Across these experiments, the integration of modalities revealed complementary facets of tumor impact: GVs enabled direct, longitudinal tracking of tumor growth; fUSI uncovered progressive disruption of sensory-evoked neural responses and large-scale network reorganization; and ULM exposed localized vascular disorganization and angiogenesis. Importantly, linking these measurements within the same animals allowed us to connect tumor expansion with simultaneous vascular remodeling and neural dysfunction, providing a systems-level view of brain–tumor interactions that would be inaccessible with any single technique.

## RESULTS

### Lentiviral delivery of gas vesicle genes enables stable ultrasound contrast in glioblastoma cells

To develop a novel imaging platform capable of tracking tumor progression and its impact on neural circuitry, we engineered a stable U87 human glioblastoma cell line, a well-established model cell line for studying glioblastoma^21–27^, to express GVs. To engineer the U87 cells, we used two previously established lentiviral vectors that together deliver the 8 genes encoding GVs derived from Anabaena flos aquae^13,28^ (**Fig. 2a**). The first vector includes genes for gas vesicle protein A (GvpA), internal ribosome entry site (IRES), and red fluorescent protein (RFP), under the control of a tetracycline response element (TRE); and reverse tetracycline-controlled transactivator (rtTA), for doxycycline-inducible expression, under the control of minimal EF1a promoter. The second vector contains genes necessary for gas vesicle formation (N, J, K, F, G, W, V) and green fluorescent protein (GFP), also under the control of TRE. The two fluorescent markers, RFP and GFP, were used to facilitate engineering.

**Figure 2:**
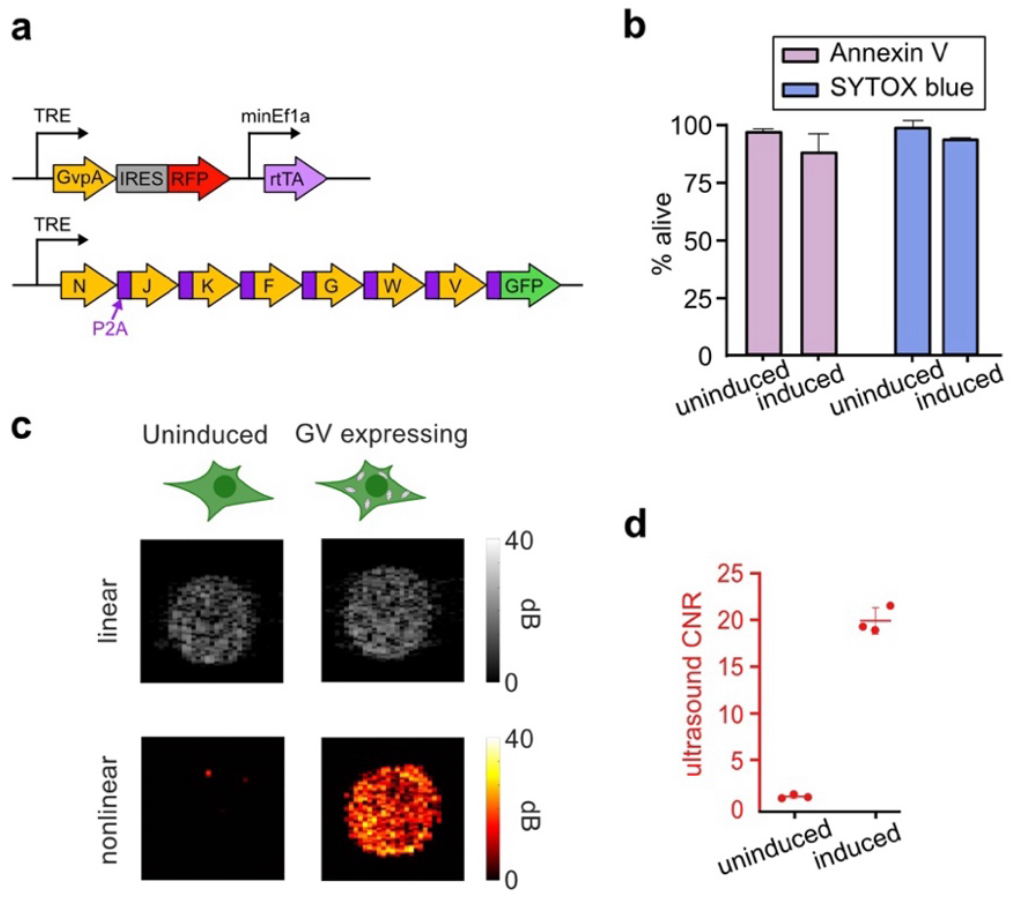
Engineering and validation of GV-expressing U87 glioblastoma cells for ultrasound imaging. **a**. Schematic representation of the genetic constructs used for engineering GV-expressing U87 glioblastoma cells. Two lentiviruses were used to deliver the 8 genes encoding GV formation, an rtTA transactivator, and two fluorescent markers. **b**. Cell viability assays using Annexin V and SYTOX Blue staining. The percentage of live cells is shown for both uninduced and doxycycline induced cells, demonstrating that the induction of GV expression does not significantly impact cell viability. **c**. Ultrasound imaging of uninduced control and dox-induced GV-expressing U87 cells. Linear (B-mode) and nonlinear (AM) ultrasound images demonstrate the enhanced contrast produced by GV expression. **d**. Contrast-to-noise ratio (CNR) of ultrasound signal from control and induced cells, highlighting the enhanced ultrasound signal from GV-expressing cells. Add error bars show mean -/+ sem.

Following lentiviral transduction, fluorescence measurements showed that 27% mean ± standard error of the mean (s.e.m) of the cell population contained both of the viral constructs. This level of positivity was substantially retained over three rounds of inductions (**Sup. Fig. 1a, b**). We found that there was no significant difference in viability between induced and uninduced cells, as measured by Annexin V and SYTOX Blue staining (**Fig. 2b**). We embedded the cells in a hydrogel phantom to enable in vitro ultrasound imaging and imaged them using amplitude modulation (AM), a method optimized to detect the nonlinear acoustic response of GVs^29^. As shown in **Fig. 2c**, both induced and uninduced samples exhibited comparable linear B-mode signals, which provide conventional ultrasound images based on tissue echogenicity and serve as a baseline measure of background cellular scattering. In contrast, nonlinear AM signals were strongly elevated in induced cells but nearly absent in uninduced controls, consistent with GV expression being required to generate nonlinear contrast. Quantification revealed a ∼20 dB increase in nonlinear signal in induced cells relative to controls (**Fig. 2d**). These experiments established our GV-expressing U87-cell line as a model glioblastoma cell line that can be robustly imaged with ultrasound, enabling our subsequent in vivo investigations.

### Trimodal ultrasound imaging reveals tumor growth, neural activity, and vascular remodeling

Having validated that our GV-expressing U87 cell line produces strong and specific ultrasound contrast in vitro, we next sought to establish an in vivo model that would allow us to longitudinally monitor tumor growth and directly link it to changes in neural activity and vascular remodeling. To this end, we implanted them into the brains of N=3 NOD scid gamma (NSG) mice to enable in vivo monitoring of tumor growth and associated functional and vascular changes in awake behaving animals (**Fig 3. a, b**). While ultrasound imaging can be performed through the intact skull of a mouse^30^, we opted to enhance image quality and signal sensitivity by performing a craniotomy and replacing the skull with a 125 μm-thick polymethylpentene (PMP) window^31^ with minimal acoustic scattering. To further support high-quality imaging in awake animals, we affixed skull head plates that allowed mice to be head-fixed within an air-lifted arena. This setup enabled the mice to move freely within a floating environment, thereby reducing anxiety and facilitating behavioral tracking under minimally stressful conditions^32^.

**Figure 3:**
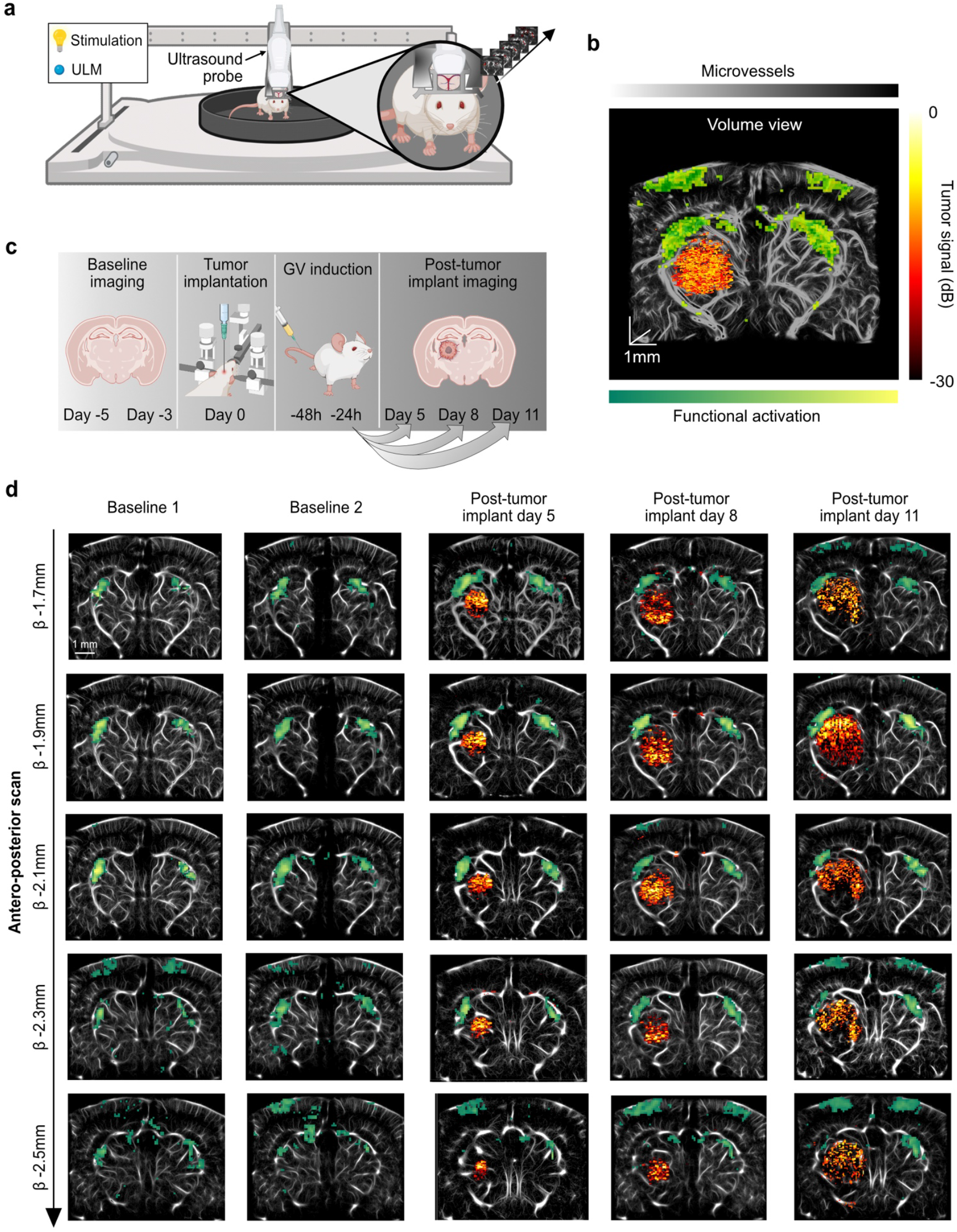
In vivo tracking of GV-expressing glioblastoma cells in mice brain using trimodal ultrasound imaging. **a**. Schematic of the experimental setup. The ultrasound probe is positioned above the animal’s head and secured to a custom head-fixation frame. The animal remains awake and can freely navigate the behavioral arena, which is supported by an air-cushioned platform to minimize stress. **b**. Experimental timeline. Baseline imaging was performed on days –5 and –3, followed by tumor cell implantation on day 0. GV expression was induced 48 and 24 hours prior to each post-implantation imaging session, conducted on days 5, 8, and 11. **c**. Representative volume renderings from trimodal imaging, showing tumor localization, vascular structure, and functional activation. Nonlinear ultrasound imaging detects GV-expressing tumor cells, ULM reveals detailed microvascular architecture, and fUSI captures visually evoked neural activity. **d**. Trimodal imaging of Mouse 1 across five anteroposterior positions (β = –1.7 mm to –2.5 mm) during baseline and post-tumor imaging sessions. Over time, the GV-expressing tumor (hot colors) becomes increasingly prominent, accompanied by alterations in the surrounding microvasculature (gray) and functional activation patterns (green). Results are representative of N=3 mice. Data from additional mice are shown in Supplementary Fig. 2.

To establish stable pre-tumor baselines and then longitudinally track tumor-induced changes in the same animals, each mouse underwent five imaging sessions: two pre-implantation baseline sessions and three follow-up scans on days 5, 8, and 11 post-implantation (**Fig. 3c**). At each time point, we performed trimodal imaging. First, we visualized the tumor using nonlinear AM, taking advantage of the acoustic signature of the GV-expressing tumor cells. Second, we recorded neural activity using fUSI under visually evoked conditions. Visual stimulation was delivered using a 5 Hz flickering blue LED, and voxel-wise correlation analysis revealed robust hemodynamic activation of the lateral geniculate nuclei (LGN). And finally, we captured high-resolution maps of the brain’s microvasculature using ULM following intravenous microbubble injection. Scans were acquired across five coronal planes per animal: β = –1.7 to –2.5 mm (0.2 mm increments) in Mouse 1, and β = –2.2 to –2.8 mm for Mice 2 and 3 (**Sup. Fig. 2**), in the anterior– posterior axis.

During baseline sessions, we validated the reproducibility of probe positioning by matching vascular landmarks across sessions. We also confirmed bilateral LGN activation in response to visual stimulation on both baseline days. Following these validations, tumor cells were stereotactically implanted into the left thalamus, 2 mm posterior to bregma. Trimodal imaging performed five days post-implantation revealed strong AM signal from the GV-expressing tumor across all imaged planes, which remained detectable throughout subsequent imaging sessions (**Fig. 3d**). Although it was found in the in vitro data that the AM signal diminished over multiple rounds of induction (**Sup. Fig. 1c, d**), this attenuation was compensated by the enhanced sensitivity of BURST^62^ imaging on the last imaging day. fUSI continued to reveal visually evoked responses, while ULM enabled longitudinal mapping of tumor-induced vascular remodeling. Representative trimodal images from all coronal planes of Mouse 1 are shown in Fig. 3d, with corresponding datasets from Mice 2 and 3 provided in **Supplementary Fig. 2**. Together these results establish that our trimodal imaging platform can robustly track tumor growth, sensory-evoked neural activity, and vascular remodeling over time within the same animals.

### Tumor growth disrupts brain structure and neural activity

Using our longitudinal dataset, we quantified tumor growth and its structural impact on surrounding brain regions (**Fig. 4a**). We observed inter-animal variability in tumor progression patterns: in Mouse 1, the tumor primarily invaded thalamic regions; in Mouse 2, it extended into both the thalamus and hippocampus; and in Mouse 3, it remained largely restricted to the hippocampus. Despite these differences, all tumors exhibited progressive expansion over time, with planar tumor areas increasing consistently across imaging sessions (**Fig. 4b**) and growth rates ranging from 0.37 to 0.53 mm^2^ per day, in line with previously reported values^33,34^. This volumetric expansion was accompanied by localized structural deformation, with noticeable displacement of the visually-activated left LGN, which lies adjacent to the implantation site and exhibited measurable shift over time.

**Figure 4:**
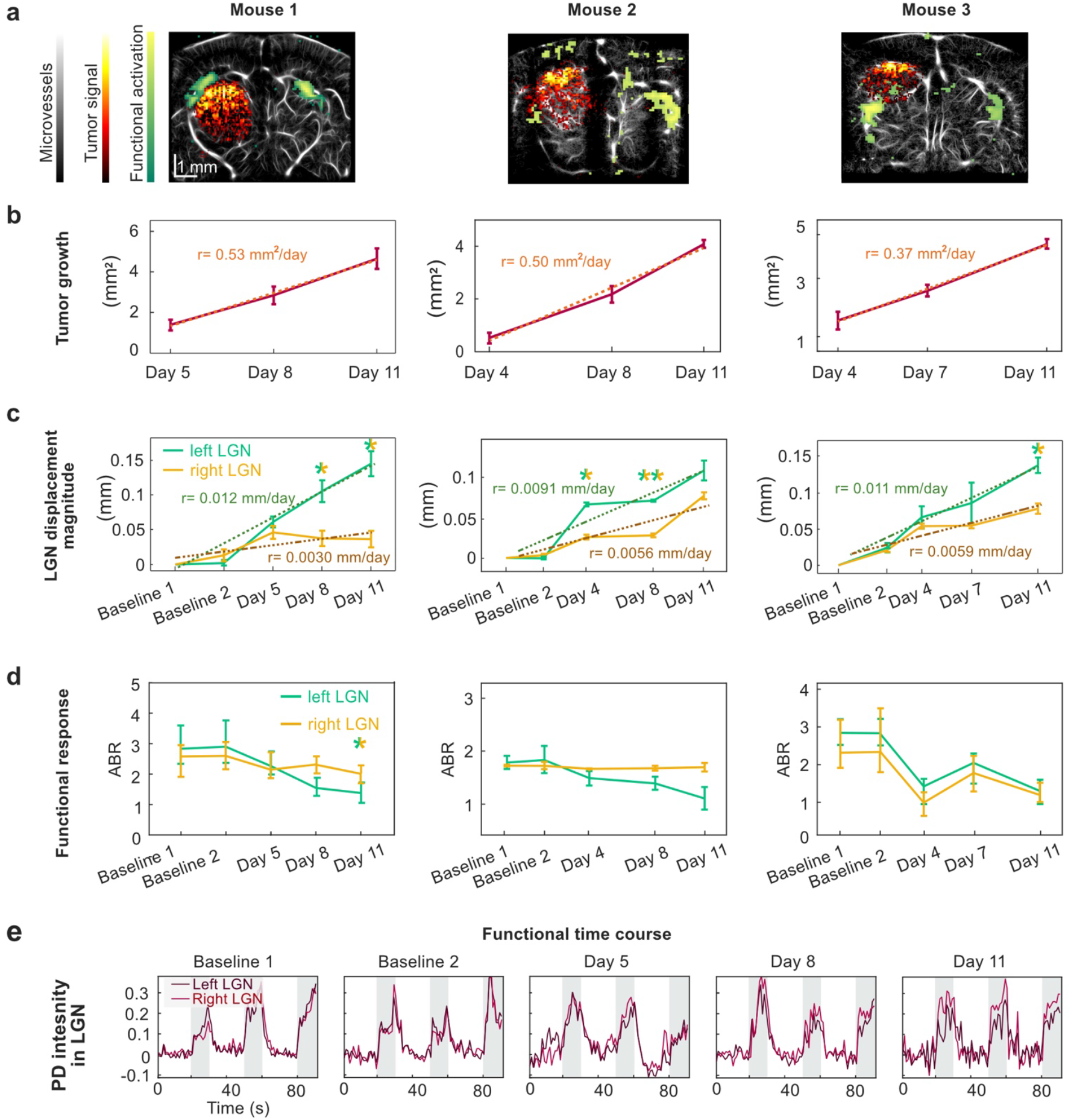
Functional and structural effects of GV-expressing glioblastoma. **a**. Trimodal imaging of mice 1, 2, and 3 at day 11 post-implantation in respective coronal plane: β-1.9mm, β-2.8mm, β-2.5mm. **b**. Planar tumor growth over time, quantified as tumor area (mm^2^) for each post-implantation imaging day. **c**. Displacement magnitude of the left and right lateral geniculate nuclei (LGNs) across all time points, including baselines. Significant interhemispheric differences were observed in: Mouse 1 at day 8 (p = 0.040) and day 11 (p = 0.029); Mouse 2 at day 4 (p = 0.036) and day 8 (p = 0.0037); Mouse 3 at day 11 (p = 0.014). **d**. Functional Activation-to-Baseline Ratio (ABR) over time. Mouse 1 exhibited a significant reduction in ABR in the left LGN by day 11 compared to the right LGN (p = 0.046). For panels (b)–(d), solid lines represent the mean per mouse across all coronal planes; error bars indicate standard error of the mean (SEM) **e**. Mean normalized hemodynamic time course from the left and right LGNs in mouse 1 across all imaging days at coronal plane β = −1.9 mm. Gray shading indicates light-on periods during visual stimulation. Detailed p-values for all comparisons are provided in Table 1.

Using optical flow–based displacement analysis^35^, we tracked LGNs’ positions over time and observed measurable displacement in all mice, albeit to varying degrees (**Fig. 4c**). The LGN on the tumor-adjacent side was consistently displaced more than the contralateral LGN, indicating that tumor mass physically deformed nearby functional brain structures. In Mice 1 and 2, displacement rates were significantly higher on the tumor-proximal side (p < 0.05), while Mouse 3 showed a milder but still detectable significant shift by day 11. These findings are consistent with prior fMRI observations in human patients, where space-occupying tumors physically displace adjacent functional brain regions^36–38^.

**Table 1:**
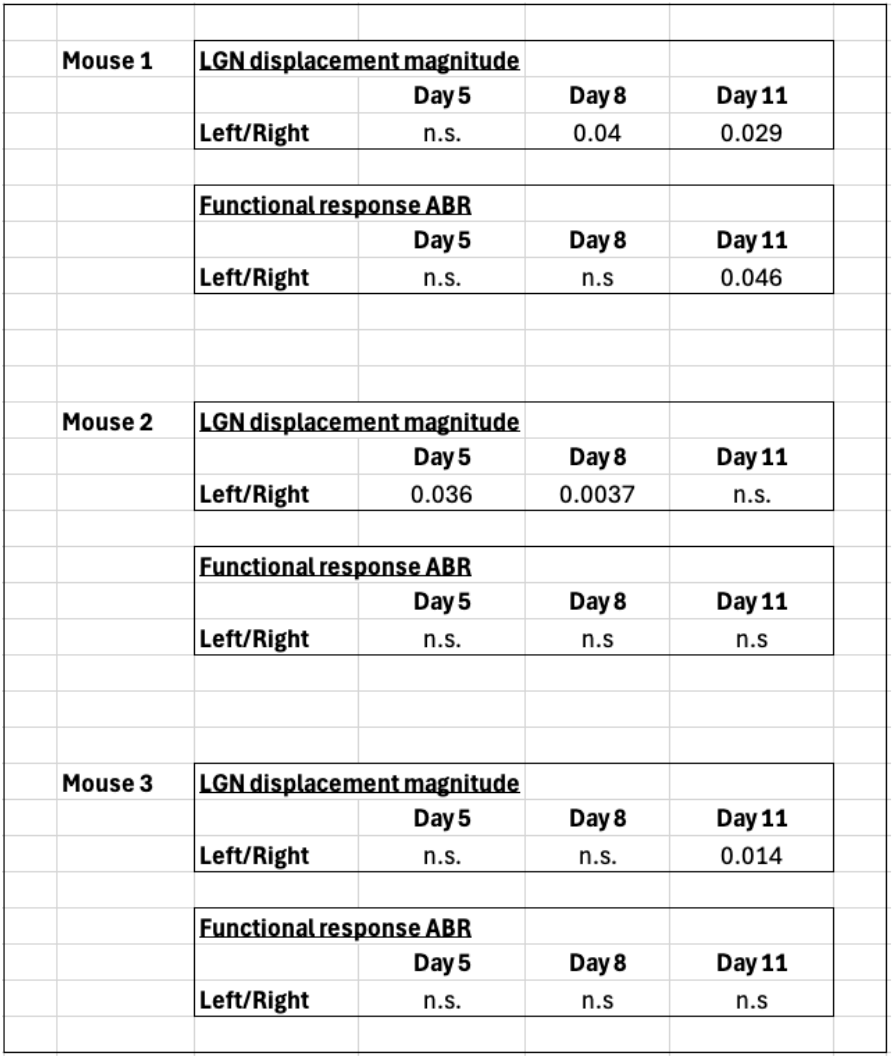
Statistical significance of the right and left LGN displacement magnitude and functional response of Mice 1, 2 and 3.

Despite the structural changes induced by tumor growth, visually evoked activation of the LGNs persisted through day 11, as measured by fUSI (**Fig. 4d, e**). However, the amplitude of these responses declined over time in the tumor-adjacent LGNs, reflected by a progressive decrease in activation-to-baseline ratios (ABR). These values were computed across all imaging planes accessible through the cranial window to ensure that the observed effects were not due to out-of-plane displacement. In Mouse 1, the ABR in the left LGN dropped significantly by day 11. Mouse 2 exhibited a similar trend, though the reduction in the left LGN was not significantly different from the contralateral side, potentially indicating more subtle or symmetric network-level adaptations. In Mouse 3, a bilateral decrease in ABR was observed, possibly reflecting systemic influences such as altered arousal state, intracranial pressure, or global hemodynamic shifts. Taken together, these results demonstrate that glioblastoma growth can induce measurable patterns of structural displacement and progressive functional impairment in both nearby and distal brain circuits. Importantly, our trimodal imaging approach provides the sensitivity and spatial resolution needed to capture this inter-animal variability, highlighting its value for detecting individualized dynamic patterns of tumor–brain interaction that would likely be missed with single-modality or population-averaged methods.

### Tumor progression disrupts local blood flow and vascular architecture

Given the variability observed across animals, we focused on Mouse 1 to illustrate the impact of glioblastoma growth on the brain’s vascular architecture, using ULM following microbubble injection (**Fig. 5**). We defined three anatomical regions of interest: the cortex, the left subcortex (adjacent to the tumor), and the right subcortex (contralateral), and quantified vessel displacement within these areas over time (**Fig. 5a–b**). Optical flow–based motion analysis^35^ in a representative coronal plane revealed progressive reorganization of the left subcortical vasculature beginning on day 5 post-implantation, while comparatively less motion was observed in the cortex and contralateral subcortex (**Fig. 5b**). Quantification from all imaged coronal planes showed that total vascular displacement in the tumor-adjacent left subcortex became significantly greater than in the right subcortex starting on day 5 (double-colored stars below the curves). Although cortical and contralateral subcortical motion also increased relative to baseline, reaching significance by day 11 (single-colored stars above the curves), the magnitude of displacement was markedly higher in the peritumoral zone. These findings support the hypothesis that tumor expansion drives localized microvascular remodeling to meet its growing metabolic demands^39^.

**Figure 5:**
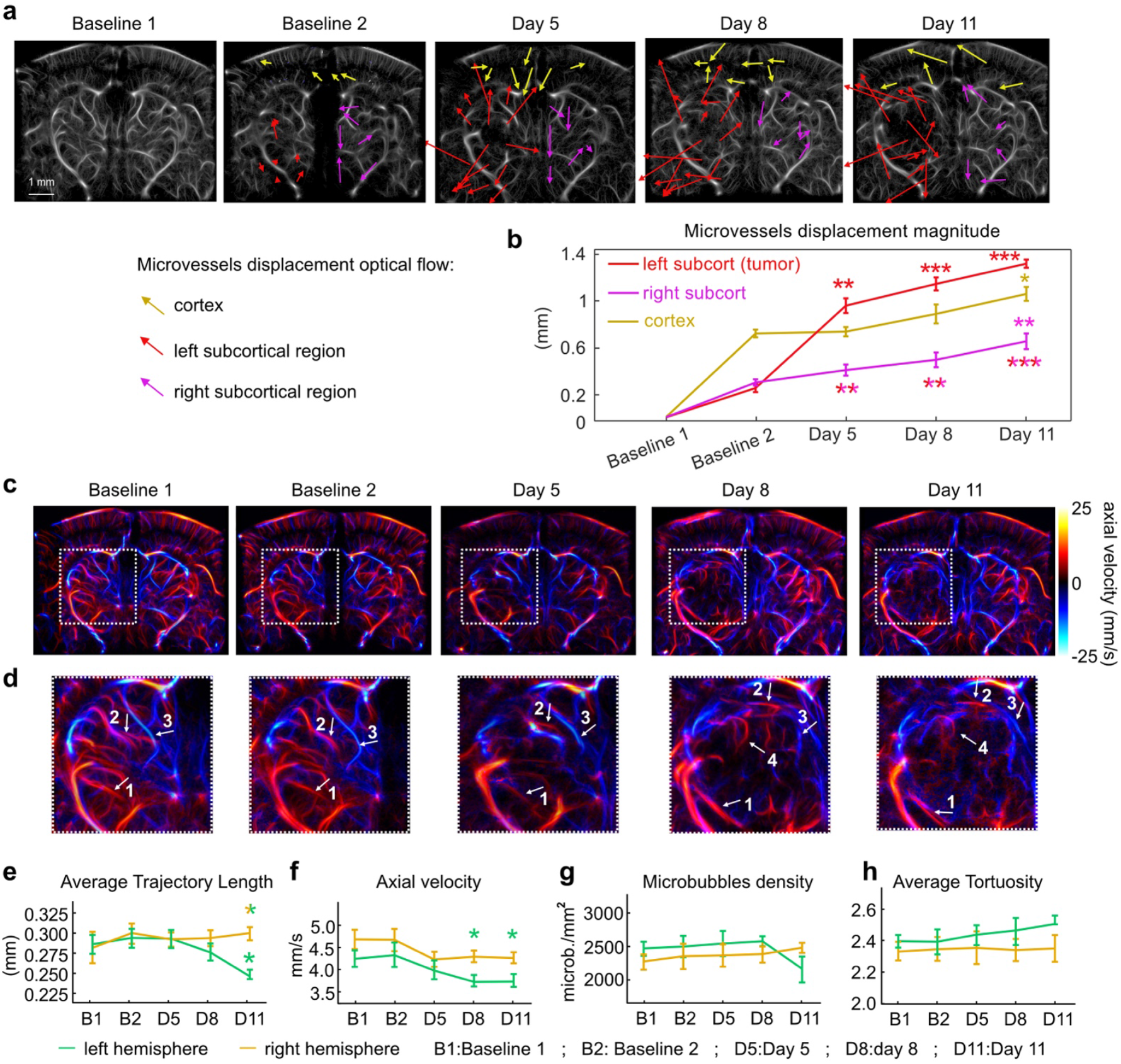
Quantification of tumor-induced microvascular alterations. **a**. Optical flow representations of microvessel displacement in the cortex (yellow), left subcortical region (red), and right subcortical region (magenta) across multiple time points. Arrows indicate the direction and magnitude of displacement. **b**. Displacement magnitude over time in each region. A significant increase in vessel motion is observed in the left subcortical region following tumor implantation, compared to both the right subcortex and cortex (double-colored stars below the curves). Within the left subcortex, displacement at all post-implantation time points is also significantly greater than baseline (single-colored stars above the curves). Error bars: SEM **c**. Two-dimensional maximum intensity projection (MIP) maps of axial microbubble velocity across the imaged volume (β = –1.7 mm to –2.5 mm, 0.2 mm increments). **d**. Magnified views of the regions outlined by white dotted boxes in (c), highlighting the trajectories of four representative vessels (white arrows 1–4) over time. **e–h**. Quantification of microvascular metrics within the tumor-adjacent (left hemisphere) region and the contralateral side. Error bars: SEM **e**. Average microbubble trajectory length between frames, **f**. Average axial flow velocity, **g**. Average microbubble density, and **h**. Average vessel tortuosity. Statistical significance: *p ≤ 0.05, **p ≤ 0.01, ***p ≤ 0.001. Detailed p-values for all comparisons are provided in Table 2.

To further explore vascular changes within the tumor core, we projected maximum intensity maps of axial microbubble velocity across timepoints (**Fig. 5c–d**). As the tumor expanded, several previously identifiable vessels were displaced from their original positions (arrows 1, 2, and 3), while new vessels emerged within the tumor region (arrow 4), including some that appeared only at later stages such as day 8. This dynamic restructuring highlights active angiogenesis and vessel recruitment within the growing tumor mass.

**Table 2:** Statistical significance of the quantification of Tumor-Induced Microvascular Alterations in Mouse 1 (related to Figure 4)

|  |  |  |  |
| --- | --- | --- | --- |
| **Microvessels displacement magnitude** |  |  |  |
|  | **D5/baseline** | **D8/baseline** | **D11/baseline** |
| **Left subcort.** | 0.002 | 0.0001 | 0.00008 |
| **Right subcort.** | n.s. | n.s. | 0.02 |
| **Cortex** | n.s. | n.s. | 0.0015 |
| **Average trajectory length** |  |  |  |
|  | **D5/baseline** | **D8/baseline** | **D11/baseline** |
| **Left hemisph.** | n.s | n.s | 0.05 |
| **Right hemisph.** | n.s | n.s | n.s |
| **Axial Velocity** |  |  |  |
|  | **D5/baseline** | **D8/baseline** | **D11/baseline** |
| **Left hemisph.** | n.s | 0.013 | 0.05 |
| **Right hemisph.** | n.s | n.s | n.s |

Quantitative assessment across all imaged coronal planes revealed several key alterations in microvascular behavior (**Fig. 5e–h**). In the left hemisphere, the average trajectory length of microbubbles between successive ultrasound frames decreased from 0.290 ± 0.124 mm at baseline to 0.276 ± 0.113 mm at day 8 and 0.246 ± 0.062 mm by day 11, a statistically significant reduction consistent with narrowing or obstruction of vascular pathways due to tumor compression or remodeling^40,41^ (**Fig. 5e**). In contrast, the right hemisphere maintained a stable microbubble trajectory (∼0.290 mm) throughout, indicating intact flow dynamics in regions unaffected by the tumor. A similar pattern was observed in axial flow velocity: in the left hemisphere, velocity declined from 4.3 ± 0.24 mm/s at baseline to 3.7 ± 0.15 mm/s by day 11, with significant reductions from baseline evident beginning at day 8 (**Fig. 5f**). The right hemisphere exhibited a slight decline, but this change did not reach statistical significance, reinforcing the localized nature of tumor-induced hemodynamic disruption. Microbubble density remained stable in both hemispheres (∼2500/mm^2^) at early timepoints but dropped to 2100/mm^2^ in the left hemisphere by day 11, indicating a reduction in the fraction of vessels remaining patent and perfused as the tumor expanded. This reduction may reflect vessel pruning, or shunting, mechanisms previously described in tumor vasculature^42^. Lastly, vessel tortuosity in the left hemisphere increased from a baseline of 2.39 ± 0.062 to 2.50 ± 0.065 by day 11 (**Fig. 5h**), reflecting the hallmark disorganization of tumor-induced angiogenesis^18,43,44^. This increased tortuosity, combined with reduced velocity and trajectory length, underscores how glioblastoma growth reshapes vascular architecture to sustain itself. Together, these findings demonstrate that glioblastoma imposes profound, region-specific changes on the brain’s vascular network, including mechanical displacement, angiogenesis, flow restriction, and eventual structural destabilization, captured in vivo with high spatial and temporal resolution using our trimodal imaging approach.

### Tumor progression reconfigures brain-wide connectivity

While our earlier findings showed that the LGNs retained stimulus-evoked responses despite mechanical displacement, we next sought to investigate how glioblastoma growth affects broader functional brain networks. To do so, we quantified resting-state functional connectivity to capture spontaneous brain activity in undisturbed animals over a continuous 10-minute period. For all three mice, we used the Paxinos and Watson^45^ mouse brain atlas to define regions of interest (ROIs) within the coronal plane where the tumor appeared largest and where multiple brain structures could be simultaneously captured (**Fig. 6a-f**). These included the cortex, corpus callosum (c-c), hippocampus (hipp), thalamus (thal), and hypothalamus (hyp-thal). Functional connectivity matrices derived from these ROIs (**Fig. 6b, d, f**) revealed at baseline states strong, bilaterally symmetric baseline networks characterized by robust intra-cortical and interhemispheric thalamic and hippocampal correlations. As the tumor progressed, however, all mice exhibited a gradual breakdown in global connectivity. By day 11, thalamic connectivity was markedly diminished in all animals, and in Mouse 1, cortical connectivity also declined, indicating broader network destabilization. Notably, in Mice 2 and 3, hypothalamic connectivity increased at later time points, possibly reflecting compensatory reorganization in subcortical networks or maladaptive hyperconnectivity, a phenomenon reported in multiple models of brain disruption^46,47^.

**Figure 6:**
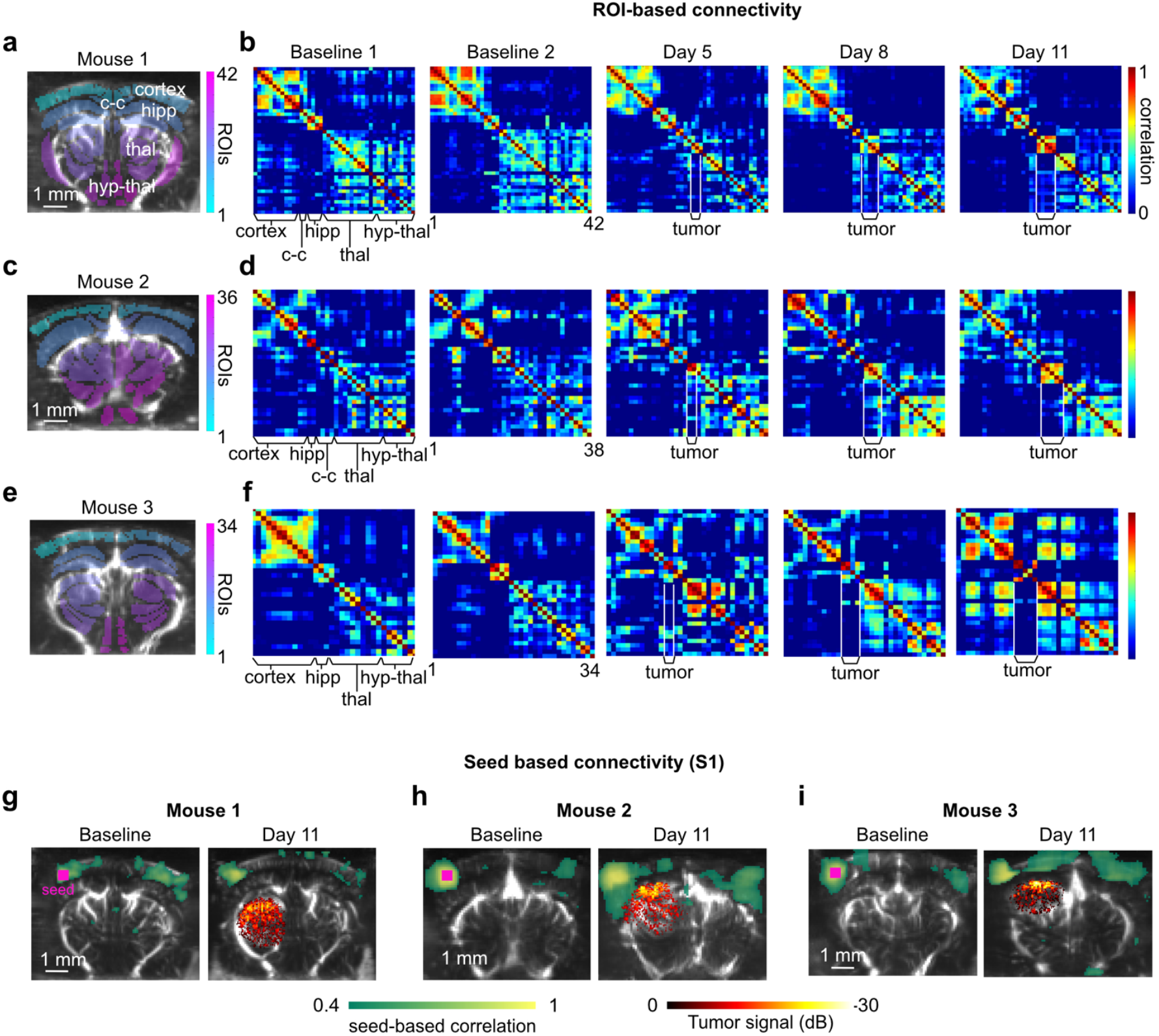
Impact of GV-expressing tumor progression on functional brain connectivity. **a, c, e:** Regions of interest (ROIs) selected for connectivity analysis in Mouse 1 (AP plane: β = –1.9 mm), Mouse 2 (β = –2.1 mm), and Mouse 3 (β = –2.5 mm). ROIs include the cortex, corpus callosum (c-c), hippocampus (hipp), thalamus (thal), and hypothalamus (hyp-thal). **b, d, f**: Functional connectivity matrices showing correlation values between ROIs at baseline (Baseline 1 and 2) and day 11 post-tumor implantation. **g–i**: Seed-based connectivity maps derived from the primary somatosensory cortex (S1) in each mouse. Correlation strength is visualized using a green colormap, overlaid with the GV-expressing tumor signal at day 11 (hot colormap) on temporally averaged power Doppler images (gray colormap) acquired during resting-state fUSI.

To further probe tumor-specific dynamics, we computed voxel-wise percent changes in CBV across time (**Sup. Videos 1 and 2**). Interestingly, the tumor region itself exhibited coherent, spatially uniform CBV fluctuations that appeared uncoupled from surrounding brain networks (e.g. T= [100s -130s] s in Sup. video 1, and T= [28s-55s] in Sup. video 2). Unlike physiological functional signals, which typically arise from coordinated activity across anatomically connected areas, these tumor-associated fluctuations lack synchronized correlations with other brain regions. This spatial and temporal isolation suggests that the tumor generates autonomous hemodynamic activity, as it has been recently observed with multiphoton microscopy^48^.

As an illustration of tumor-induced network reorganization, we performed seed-based connectivity analysis using the primary somatosensory cortex (S1) as a seed region (**Fig. 6, g-i**). At baseline, all animals exhibited strong bilateral and symmetric connectivity between S1 and the contralateral cortex, consistent with known interhemispheric coordination of sensory networks. However, by day 11 post-implantation, connectivity patterns diverged markedly across animals. In Mouse 1, where the tumor expanded primarily within thalamic regions, S1 connectivity decreased significantly, with weakened interhemispheric correlations and reduced integration with surrounding cortical territories. In contrast, Mice 2 and 3 — both of which developed tumors that extended into the hippocampus — showed expanded S1 connectivity, with the seed-based network encompassing broader ipsilateral and contralateral cortical areas. These distinct trajectories highlight how localized tumor growth can have variable and widespread effects on cortical networks, and suggest that tumors may induce both degenerative and compensatory remodeling of functional connectivity, depending on their anatomical location and local circuit interactions^49–51^. Taken together, these results demonstrate the far-reaching and dynamic effects of glioblastoma progression on functional brain architecture and its variability between subjects. Beyond local displacement and impairment, tumors reshape global network topology, giving rise to aberrant, tumor-centered hubs and the collapse of connectivity patterns. These findings underscore the importance of whole-brain, longitudinal imaging approaches in capturing the distributed and evolving nature of tumor-induced functional disruption.

## DISCUSSION

Our study introduces and validates a trimodal ultrasound imaging platform that leverages genetically encoded tumor labels, hemodynamic functional imaging, and super-resolved vascular mapping to non-invasively monitor glioblastoma progression and its impact on brain structure and function. This approach allows for high-resolution, longitudinal observation of tumor-brain interactions. Our analysis revealed significant displacement of functional regions such as the LGN in the tumor hemisphere, and concurrent ULM imaging showed extensive vascular remodeling, including increased vessel tortuosity and tumor-induced angiogenesis. Despite the physical distortion, we observed persistent, but attenuated, visually evoked activity in the LGNs, highlighting a degree of functional resilience, aligning with previous observations where displaced or lesioned regions can retain activity through plasticity or network compensation^52–54^. Quantitative analyses of ULM-derived flow maps revealed progressive shortening of microbubble trajectories, reduced flow velocity, and increased vessel tortuosity in the tumor hemisphere, consistent with glioblastoma-driven vascular constriction and disorganized angiogenesis^18–20^. Through whole-brain functional connectivity analyses, we consistently observed a progressive remodeling of inter-regional connectivity, even in regions clearly outside the tumor growth area.

While the goal of this study was to introduce and validate our multimodal platform, there are several important directions in which the approach can be expanded. One set of opportunities involves testing variables that would further probe the scalability of the system. For example, we focused here on a single tumor model, U87 glioblastoma, which is widely used but does not fully capture the invasive properties of more diffuse models such as diffuse intrinsic pontine glioma or metastatic cascade. Applying the platform to smaller or diffused tumor models would also help determine its sensitivity. In addition, our work was limited to a single implantation site in the left thalamus and to one evoked sensory paradigm, visual stimulation. Exploring other implantation sites across brain regions, as well as incorporating additional paradigms such as auditory or somatosensory stimulation, could provide a more comprehensive view of tumor-induced dysfunction. Furthermore, the platform opens the door to a wide range of new applications beyond tumor characterization alone. It can be combined with therapy to investigate treatment effects on both tumors and surrounding neural circuits in vivo. It also provides a foundation for integrating molecular-scale readouts by pairing fUSI with genetically encoded calcium or voltage indicators, offering a unique window into cell-type–specific or neurotransmitter-specific dynamics alongside whole-brain hemodynamics^55^. Finally, while our sample size was limited, the reproducible trends suggest that future studies can test the generality of these findings and determine whether tumor-related brain activity changes are conserved across subjects.

Together, these findings emphasize the complex interplay between structural disruption, vascular remodeling, and functional network reorganization. The ability of our platform to capture all three dimensions of tumor progression in vivo in awake animals represents a major step forward in studying neuro-oncological processes with spatiotemporal precision and developing improved treatments.

## METHODS

### Statistical analysis

All statistical comparisons were performed using two-tailed unpaired Student’s t-tests unless otherwise specified. Results are reported as mean ± standard error of the mean (s.e.m.), and significance thresholds were set at p < 0.05. Exact p-values and sample sizes (n) are provided in figure legends or in supplemental tables. Statistical analyses were conducted in MATLAB (MathWorks) and Prism (GraphPad).

### Generation of Gas Vesicle (GV) – Expressing Glioblastoma Cells

DMEM media was prepared by supplementing high-glucose DMEM with GlutaMAX, 1 mM sodium pyruvate (Gibco, Catalog #10569), 20 mM HEPES (Cytvia, Catalog #SH30237), 1X MEM-Non-Essential Amino Acids (0.1 mM, Gibco #11140), and 10% fetal bovine serum (bio-techne, Catalog # S10350). HEK293T cells (ATCC, CRL-3216), passaged no more than 20 times, were seeded in 10 cm plates and grown in DMEM+ until they reached 80 - 90% confluence. A mixture of 22 µg pLV (plasmid with the gene of interest), 22 µg pCMVR8.74 (Addgene #22036), and 4.5 µg pMD2.G (Addgene #12259) was prepared. These plasmids were combined with polyethylenimine (PEI, Polysciences) at a PEI to DNA mass ratio of 2.8:1, incubated at room temperature for 12 minutes, and then added to the cells. Cells were incubated at 37°C and 5% CO2 for 12 hours. After this period, the transfection media was replaced with 10 mL of fresh DMEM+ supplemented with sodium butyrate (10 mM) for 8 hours. The media was then replaced with 10 mL of fresh DMEM+. After 48 hours, the spent media was collected and centrifuged at 500x g for 10 minutes to remove debris. The viral supernatant was stored at 4°C and used for transduction within 24 hours of collection, as described below.

U87 cells (ATCC) were seeded in DMEM+ media in 6-well plates 24 hours before transduction and grown to 70 - 90% confluence. 500 uL of each lentivirus was warmed up to 37°C and added to each well with 10 µg/mL of polybrene (Millipore Sigma, Catalog #TR1003). Cell culture plates were spun at 800 xg for 50 minutes at 35°C, and incubated at 37°C for 16-20 hours. The viral media was then removed, and the cells were passaged for downstream applications. Cells were induced with doxycycline (1 ug/mL) for 72 hours prior to measurements. Transduction efficiency was confirmed with flow cytometry (MACSQuant Vyb) after one, two and three rounds of induction. Cell viability post-transduction was assessed using Annexin V (Invitrogen, Catalog #A35136) and SYTOX Blue (Invitrogen, Catalog #S34857) staining following manufacturer’s protocols. GV-expressing cells were embedded in agarose phantoms soaked in 1X PBS for in vitro ultrasound imaging (**Sup. Fig.1.c,d**).

### GV preparation for in-vivo experiments

GV-expressing U87 cells were cultured without doxycycline induction for at least 2 passages before in vivo implantation. On the day of implantation in the mouse brain, cells were dissociated using 0.25% Trypsin/EDTA (Corning 25-053-CI), centrifuged at 500 xg for 5 minutes, and resuspended in Phosphate-Buffered Saline (PBS) at a concentration of around 60 million cells per milliliter. The cells were then stored for up to 2 hours on ice before injection.

### Craniotomy and implantation of a sonotransparent skull window

All animal procedures were conducted in compliance with the Institutional Animal Care and Use Committee guidelines of the California Institute of Technology (Protocol 1761). Immunocompromised NSG mice (both male and female, aged 6–14 weeks; Jackson Laboratory) were used in this study. Randomization and blinding were not applicable due to the experimental design. Prior to surgery, mice were weighed and anesthetized with 4% isoflurane in room air and maintained under 1–2% isoflurane throughout the procedure. Body temperature was kept at 37 °C using a heating pad. For perioperative analgesia, 150 µL of ketoprofen (1 mg/mL), 50 µL of sustained-release buprenorphine (Ethiqa XR, 1.3 mg/mL), and 50 µL of bupivacaine (0.5 mg/mL) were administered subcutaneously, with bupivacaine injected directly at the incision site. The scalp was shaved and cleaned with chlorhexidine, then incised and retracted to expose the skull. A craniotomy was performed using a micro drill with a steel burr, removing a skull segment spanning from anterior-posterior (AP) coordinates -0.5 mm to AP -3.5 mm and medial-lateral (ML) coordinates -2.5 mm to ML +2.5 mm relative to bregma. Care was taken to preserve the integrity of the dura. The excised skull was replaced with a sonotransparent polymethylpentene (TPX) window (125 µm thick) of equivalent size. This window was affixed to the skull using a UV-curable composite (Tetric EvoFlow) and reinforced with dental cement (Metabond, Patterson Dental). To facilitate imaging in a head-fixed setup, a custom-designed headplate was attached to the cement base anchoring the sonotransparent window. This headplate was specifically designed to interface with the Mobile HomeCage system, enabling stable head fixation while permitting free movement of the animal within the behavioral cage. Postoperative recovery was rapid (3 days), with animals exhibiting normal behavior, including being bright, alert, and responsive. Animals were closely monitored to ensure recovery and overall well-being.

### Implantation and induction of GV-expression glioblastoma cells

Following the two baseline recording sessions, GV-expressing U87 glioblastoma cells were stereotactically implanted into the brains of the mice. Mice were weighed and anesthetized with 4% isoflurane in room air, with anesthesia maintained at 1– 2% isoflurane throughout the procedure. Body temperature was regulated at 37 °C using a heating pad. Analgesia was provided via a subcutaneous injection of 150 µL ketoprofen (1 mg/mL). A small access hole was created in the sonotransparent window using a micro drill at coordinates anterior-posterior (AP) -4 mm and medial-lateral (ML) +1.5 mm relative to bregma. A 33G needle (World Precision Instruments) attached to a microliter syringe (Hamilton) was positioned at a 45° angle and inserted 4 mm into the brain to reach the target tumor implantation site: Anterior-Posterior (AP) -2 mm, medial-lateral (ML) +1.5 mm, and dorsal-ventral (DV) -3.5 mm. Tumor cells were injected at a flow rate of 5–7 nL/s using a micro syringe pump (World Precision Instruments). A total volume of 1.5–3 uL, containing 10^5^ cells, was delivered. After waiting 5 minutes to allow cell stabilization, the needle was gently retracted, and the access hole was sealed with a UV-curable composite (Tetric EvoFlow). Postoperative recovery was rapid (3 days), with animals displaying normal behavior, including being bright, alert, and responsive. Animals were closely monitored post-surgery to ensure full recovery and continued well-being. Cells were induced to express GVs 48 and 24 hours prior to imaging sessions by intraperitoneal injection of doxycycline (0.1 mg/mL). Mice were allowed to recover and monitored daily to ensure health and stability.

### Ultrasound Imaging Setup

Trimodal imaging was conducted using a Verasonics Vantage programmable ultrasound system, controlled via custom MATLAB (MathWorks, USA) scripts. The system was equipped with an L22-14vX 128-element linear array transducer, featuring a 0.1 mm pitch, an elevation focus of 8 mm, an elevation aperture of 1.5 mm, and a center frequency of 15.25 MHz with a -6 dB bandwidth of 67%. To minimize stress and ensure physiological activity during imaging, mice were trained to acclimate to the head-fixed Mobile HomeCage system (Neurotar, Finland) prior to the first imaging session. This training involved placing mice in the air-cushion-supported behavioral cage for 1 to 3 hours daily over a period of five days. This habituation period allowed the mice to adapt to the setup and reduced stress during subsequent imaging sessions. On imaging days, mice were briefly anesthetized with isoflurane (<1min) to facilitate secure positioning in the head-fixed setup. Once head-fixed, isoflurane was discontinued to allow the mice to regain consciousness. Imaging sessions began 10 minutes after cessation of isoflurane to ensure its complete clearance from the system, minimizing potential confounds such as vasodilation. Trimodal imaging was performed sequentially, beginning with tumor visualization using AM imaging, followed by functional recordings of brain activity (visual-evoked and functional connectivity) using fUSI, and finally with microvascular mapping using ULM. Trimodal imaging was performed on baseline days -5 and -3 pre-implantation and at days 5, 8, and 11 post-implantation, enabling longitudinal tracking of tumor-induced structural and functional changes.

### Functional Ultrasound Imaging

Functional ultrasound imaging (fUSI) visualizes neural activity by mapping local changes in cerebral blood volume (CBV). CBV variations are tightly linked to neuronal activity through the neurovascular coupling^56^ and are evaluated by calculated power Doppler variations in the brain^57^. Each power Doppler image was obtained from the accumulation of 200 compounded frames acquired at 500 Hz frame rate. Each compounded frame was created using 14 tilted plane waves (-14° to 14° by step of 2°). We used a pulse repetition frequency (PRF) of 7000 Hz. fUSI images were repeated every second. Each block of 200 images was processed using a SVD clutter filter^58^ to separate tissue signal from blood signal to obtain a final power Doppler image exhibiting CBV in the whole imaging plane. Functional ultrasound imaging movies were created by repeating the acquisition of power Doppler images at a framerate of one power Doppler image per second.

#### Evoked-activation maps following visual stimulation

We used a passive visual simulation block task designed to activate the visual system. Visual stimuli were delivered using a blue LED (450 nm wavelength) positioned at 8 cm in front of the head of the mice. Stimulation runs consisted of periodic flickering of the blue LED (flickering rate: 5 Hz) using the following parameters: 30 s dark, followed by 30 s of light flickering repeated three times for a total duration of 180 s. At this distance, the light luminance was of 14 lux when the light was on and ∼0.05 lux when the light was off. Correlation maps were computed individually from the normalized correlation between each voxel temporal signal with the different stimulus patterns (Pearson’s product moment) using MATLAB (MathWorks). We chose a level of significance of z > 3.1 by applying the Fisher’s transform (P < 0.001, one-tailed test), which corresponds to r > 0.193. This allowed us to assess the statistical significance between the hemodynamic response and the stimulation structure for each voxel. For all the scanned coronal brain planes, we identified voxels within the lateral geniculate nucleus (LGN) activated during these visual stimulation.

#### Functional connectivity analysis

To study functional connectivity, we recorded the functional activity of the head-fixed awake mice during 10 min per imaging plane, without any external stimulation (while the mice were kept in a light-and-noise isolated chamber). We used established analysis to process our functional connectivity recordings^59^. We first removed motion artifacts from each acquisition by thresholding on Doppler signals chosen at +/- 10% of the median value, and of all remaining frames we only kept frames that formed a successive uninterrupted period of 30 s, to avoid fragmented or small individual periods. We then applied a low-pass filter at 0.2 Hz to remove high frequency signals while preserving the resting-state frequency band. The signal was then detrended with a polynomial fit of order 4 to remove low frequency fluctuations. Finally, singular value decomposition of the data was performed. Since the first spatial eigenvector is coupled to the whole brain tissue, it was systematically removed. Although this form of global signal regression can lead to anti-correlation values between structures that may be difficult to interpret, it was preferred to avoid any remaining global motion signal that can affect the whole brain tissue and induce artificially strong correlations. To build functional connectivity matrices [M], we used the time course of the CBV signal extracted from N different ROIs defined from the Paxinos Atlas^45^. We then determined the Pearson correlation between each filtered ROIs’ signal and report the correlation coefficients on a N × N matrix. At the intersection of the ith line and jth column, the value of the cell M_i,j_ of the matrix M is the correlation coefficient between the region i and the region j. Seed-based maps were also calculated from the Pearson correlation over time of the time course of the CBV signal extracted from a small selected ROI (the seed region as defined by the operator), with signals from all the other pixels of the brain.

### Ultrasound Localization Microscopy (ULM)

Microvascular architecture was mapped using Ultrasound Localization Microscopy (ULM). While under anesthesia, the animal was injected with 50 ul of Activated DEFINITY® (Lantheus, Billerica, MA, USA) via a tail vein catheter diluted at 1/10 of the original concentration with sterile saline. The craniotomized brain is imaged using ultrafast ultrasound at a compounded imaging rate of 5000 Hz (6 angles equally spaced from -5 to 5 degrees) during 0.5 s. A pause is implemented for saving and this sequence is repeated 250 times to yield 100k images. The transmitted pulse is centered at 15.625MHz and spans two cycles. The RF data is sampled at 100% and then beamformed using the delay-and-sum beamformer provided by the Vantage API (Verasonics, Kirkland, WA, USA). The beamformed data is cropped to reduce the region of imaging to the brain only, then filtered using a 3^rd^ order Butterworth bandpass filter in between 15 and 249Hz, followed by an additional spatio-temporal clutter filter based on ingular Value Decomposition^58^ (SVD), removing the 2 highest energy singular vectors. An adaptive TGC is applied to enhance microbubble signal based on filtered Power Doppler data (SVD-filtered, removing 4 of the highest energy singular vectors). The 1-lag auto-correlation further enhances the microbubble signal. Localization and tracking of microbubbles are then performed using a radial symmetry algorithm^60^ and a Munkres assignment pairing algorithm for tracking.

### Quantification of ULM Trajectory Features

To quantify local vascular properties from ULM data, we analyzed individual microbubble trajectories extracted from super-resolved ULM acquisitions. Regions of interest (ROIs) were manually drawn over both the tumor core and matched contralateral brain regions using Power Doppler images as anatomical references. Only trajectories that intersected each ROI were included in the analysis. For each included trajectory, we calculated four metrics: 1-Axial velocity: Mean axial (depth-wise) velocity along the trajectory, as reported by ULM tracking. 2- Trajectory length: The linear distance between the first and last coordinates of the trajectory, representing the straight-line displacement. 3- Tortuosity: Computed as the ratio of the total path length to the straight-line distance between trajectory endpoints, capturing deviations from linearity. 3- Microbubble density: Estimated as the number of trajectories intersecting the ROI divided by the ROI area (in pixels), serving as a proxy for local perfusion. This analysis was performed independently for the tumor and contralateral ROIs in each imaging session. Trajectories with missing or invalid points were excluded, and tortuosity values were discarded if total trajectory length was zero.

### Nonlinear Ultrasound Imaging of GV-Expressing Glioblastoma

To visualize GV-expressing glioblastoma cells, we leveraged the nonlinear acoustic scattering properties of gas vesicles (GVs), which arise from reversible shell buckling under pressure. In vitro imaging was performed using nonlinear X-Wave Amplitude Modulation (xAM) ultrasound imaging^29^, a plane-wave–based technique that creates localized amplitude modulation at the intersection of two angled wavefronts. This geometry confines nonlinear contrast generation to the focal region while minimizing background accumulation from propagation-induced nonlinearities. In vivo imaging was performed through the 125 μm-thick polymethylpentene cranial window of the mice using parabolic amplitude modulation (pAM)^61^ which delivers focused ultrasound through the skull implant with greater beam coherence than plane-wave methods. The pAM sequence consisted of three successive pulses at amplitudes 1, ½, and ½, transmitted using distinct transducer element patterns. Nonlinear signals were isolated by subtracting the combined half-amplitude responses from the full-amplitude signal. For the final in vivo imaging session (day 11), we employed BURST imaging^62^ to enhance sensitivity. BURST combines amplitude modulation with temporal signal unmixing to maximize sensitivity to nonlinear scatterers. Given the observed signal reduction after repeated doxycycline induction in vitro (**Sup. Fig. 1.c,d**), BURST was chosen to improve detectability despite potential GV collapse, which was acceptable on the final day of imaging. This sequential multimodal approach enabled robust and high-contrast visualization of GV-expressing glioblastoma across experimental stages and biological conditions.

### Tumor growth quantification

To quantify tumor growth over time, we analyzed the nonlinear AM signal of GV-expressing cells across imaging sessions at days 5, 8, and 11 post-implantation. AM images were first normalized by their maximum value, and voxels were classified as tumor-positive if their intensity exceeded 2.58 standard deviations above the mean, corresponding to a z-score > 2.58 (p < 0.01). Because thresholded voxels did not always form a contiguous mass, we generated a continuous tumor mask by computing an alpha shape around the positive voxels and rasterizing the enclosed volume (MATLAB alphaShape, inShape), which bridged small gaps while preserving non-convex morphology. Tumor area was quantified for each imaged plane by converting the number of positive pixels to physical units (mm^2^) based on voxel size. This procedure was repeated across all coronal planes, and mean tumor areas with standard error of the mean (s.e.m.) were reported across planes for each day.

### Optical Flow–Based Displacement Analysis

We performed optical flow–based displacement analysis to quantify tumor-induced deformation over time of the LGN (**Fig. 4**) and of the cerebral microvessels (Fig. 5), following methods adapted from Afrashteh et al.^35^. Activation maps (for the LGN) or ULM maps (for the microvessels) from five timepoints (baseline 1, baseline 2, day 5, day 8, and day 11) were stacked into a 3D array. To analyze spatial displacement patterns, we applied the Farnebäck dense optical flow algorithm (opticalFlowFarneback) in MATLAB. Optical flow fields were computed between the baseline reference and each subsequent timepoint, yielding pixel-wise motion vectors representing displacement over time. For Fig. 5a, optical flow fields were visualized by overlaying vector plots onto the corresponding ULM maps. Each ULM map was intensity-normalized (cube-root transformed to compress dynamic range) and displayed in grayscale. Flow vectors were then plotted using a decimation factor of [80 80] to reduce visual clutter and a scale factor of 10 to enhance visibility, resulting in a qualitative representation of the magnitude and direction of tissue deformation across hemispheres.

### Functional Response Comparison: Activation-to-Baseline Ratio (ABR) Analysis

To assess changes in visually evoked responses over time, we computed activation-to-baseline ratios (ABRs) from normalized hemodynamic time courses extracted from the lateral geniculate nucleus (LGN). For each mouse and imaging day (baseline 1, baseline 2, day 5, day 8, and day 11), time courses were normalized to z-scores and grouped into a matrix representing each session. For each session, we computed the ABR as the difference between the mean signal during stimulation and the mean signal during baseline, normalized by the standard deviation of the baseline signal. ABRs were calculated independently for each imaging plane (e.g., β = – 1.9 mm, –2.7 mm, etc.). Values were then pooled across planes to compute mean and standard deviation at each timepoint.

### Figures

Diagrams were generated using BioRender.

## FUNDING

Human Frontier Science Program Cross-Disciplinary Fellowship (LT000217/2020-C) : CR

National Institutes of Health (R01-EB018975, R01NS120828): MGS

The Mark Foundation: MGS

National Science Foundation Graduate Research Fellowship Program fellowship: SS

Howard Hughes Medical Institute: MGS

## AUTHOR CONTRIBUTIONS

Conceptualization: CR, MGS

Genetic design and lentiviral delivery: SS

Surgeries and in vivo experiments: CR

ULM experiment design and processing: BH

Ultrasound data processing and analysis: CR

Supervision of the research: MGS

Writing: CR, SS, BH, MGS

## DATA AND MATERIALS AVAILABILITY

Key data used in this paper will be stored on CaltechDATA (https://data.caltech.edu/), or a similar data repository. DOI accession number will be generated upon acceptance of paper. Code used to generate key figures and results will be posted to a publicly accessible GitHub repository and an archived version will be stored on Zenodo or similar archivable code repository. DOI accession number will be generated upon acceptance of paper.

## SUPPLEMENTARY MATERIALS

**Supplementary Figure 1.**
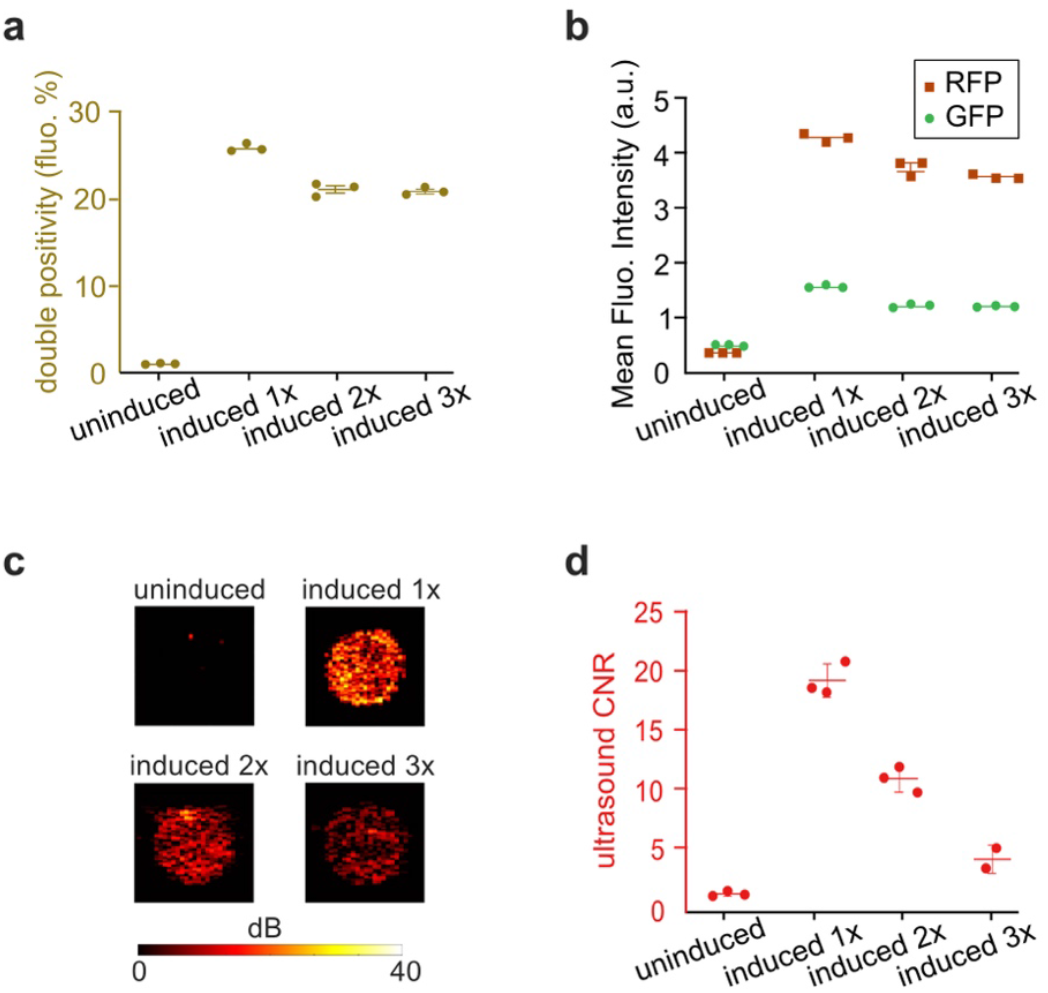
Fluorescence and ultrasound characterization of GV expression in engineered U87 glioblastoma cells across induction cycles. **a**. Percentage of double-positive cells (GFP and RFP) as measured by flow cytometry for uninduced and doxycycline-induced cultures (1×, 2×, or 3× inductions). Each dot represents an independent biological replicate. **b**. Mean fluorescence intensity (a.u.) of GFP and RFP reporters in the same conditions, showing stable expression across repeated inductions. **c**. Representative nonlinear X-Wave ultrasound images of cell pellets under each condition. Induction increases GV-associated signal up to 1×, with subsequent reductions in signal at 2× and 3×. **d**. Quantification of ultrasound contrast-to-noise ratio (CNR) reveals maximal signal after a single induction, followed by progressive decreases at 2× and 3×. Data shown as mean ± SEM.

**Supplementary Figure 2.**
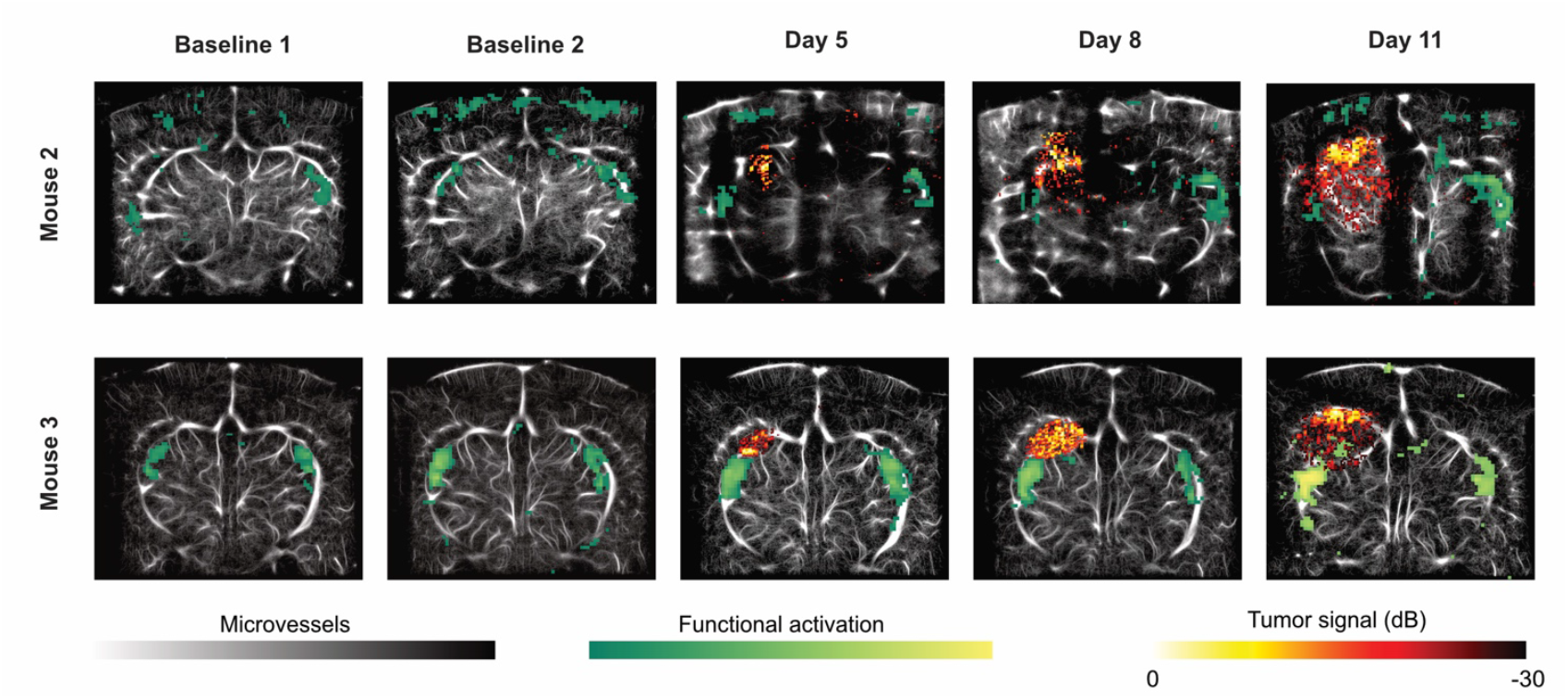
Longitudinal trimodal ultrasound imaging of GV-expressing glioblastoma in Mice 2 and 3. Representative coronal trimodal images from β = –2.8 mm (Mouse 2, top row) and β = –2.5 mm (Mouse 3, bottom row) across five imaging sessions: two pre-implantation baselines and three post-implantation days (5, 8, and 11). Each panel shows: the microvasculature from ultrafast localization microscopy (ULM; grayscale), the visually evoked functional activation from fUSI (green to yellow heatmap), and the tumor signal from nonlinear acoustic imaging (xAM or pAM; red to yellow heatmap, in dB). In Mouse 2, air bubbles trapped beneath the sonotransparent skull implant disrupted microbubble tracking for ULM on Days 5 and 8, leading to degraded microvascular contrast in these sessions.

https://data.caltech.edu/records/9vtya-smf11?token=eyJhbGciOiJIUzUxMiJ9.eyJpZCI6IjI2Y2ZiZDg5LTIyNDAtNDg3ZC1iNjFmLWUzNDEzZTIyODBlZCIsImRhdGEiOnt9LCJyYW5kb20iOiIzMmYwYzVjMzM3OTgxNDMzZTExYTlkYjM2NGVkOTdhMyJ9.e8BZehiG3wD19JGJnMv91c2hpQZ3d46xHFKWsv752cq5b83KIRS4x5qpF6at5hkpD0GIC9FCLorDoCc8xC97og

**Supplemental Video 1**. Voxel-wise CBV fluctuations in Mouse 1 at day 8 post-tumor implantation. Time-lapse visualization of voxel-wise percent changes in cerebral blood volume (CBV) during the first 140 seconds of the 10-minute resting-state fUSI acquisition in AP coronal plane: β = –2.1 mm.

https://data.caltech.edu/records/qtcz6-gnk52?token=eyJhbGciOiJIUzUxMiJ9.eyJpZCI6IjRiNWNmMTJhLTdjZDUtNDIzOC1iMmYzLWIxYWIxN2M1YzIyZSIsImRhdGEiOnt9LCJyYW5kb20iOiIyODc1ZTMzOWY2Yzc0NmY4MTg0YjdhNDBjMzkzNDYwYyJ9.KtK-HLqhHQKsa5vWRh1oFSrpbwN67tLMy2-hgzUHBMnDjWn4AdwSwWs3w9Mf2bLc9Grl4ZP7LVeCMDc_Vsug1g

**Supplemental Video 2**. Voxel-wise CBV fluctuations in Mouse 1 at day 11 post-tumor implantation. Time-lapse visualization of voxel-wise percent changes in cerebral blood volume (CBV) during the first 240 seconds of the 10-minute resting-state fUSI acquisition in AP coronal plane: β = –2.1 mm.

